# The dominant axes of lifetime behavioral variation in honey bees

**DOI:** 10.1101/2021.04.15.440020

**Authors:** Michael L Smith, Jacob D Davidson, Benjamin Wild, David M Dormagen, Tim Landgraf, Iain D Couzin

## Abstract

Insect colonies are decentralized systems that employ task allocation, whereby individuals undertake different roles to fulfil colony needs, such as honey bee “nurses”, “nest workers”, and “foragers”. However, the extent to which individuals can be well-classified by discrete “roles”, how they change behavior from day-to-day, over entire lifetimes, and with environmental conditions, is poorly understood. Using long-term automated tracking of over 4,200 individually-identified bees *Apis mellifera*, we use behavioral metrics to quantify and compare behavior. We show that individuals exhibit behavioral variation along two dominant axes that represent nest substrate use and movement within the nest. Across lifetimes, we find that individuals differ in foraging onset, and that certain bees exhibit lifelong consistencies in their movement patterns. Furthermore, we examine a period of sudden nectar availability where the honey stores tripled over 6 days, and see that the colony exhibits a distributed shift in activity that did not require a large-scale colony reorganization. Our quantitative approach shows how collective units differ over days and lifetimes, and how sources of variation and variability contribute to the colony’s robust yet flexible response.

## Introduction

Social insect colonies are groups of individual organisms that form a cooperative unit to propagate their genes (***Seeley, 1989a; Smith and Szathmary, 1995***). To survive, grow, and reproduce, these superorganisms must navigate all the same biotic and abiotic challenges as unicellular and multicellular organisms, but with coordination occurring at the level of independent individuals (***Hölldobler et al., 2009***). Lacking centralized control, insect colonies are known to self-organize tasks among workers (***Oster and Wilson, 1978; Beshers and Fewell, 2001***), whether physiologically (***Porter and Tschinkel, 1985; Robinson et al., 1989, 2009a***), spatially (***Mersch et al., 2013***), and/or temporally (***Seeley, 1982***).

Task allocation is widespread among ants, bees, termites, and wasps (***Oster and Wilson, 1978; Hölldobler and Wilson, 1990; Bourke and Franks, 1995***). Physical castes in ants, for example, demonstrate the benefit of having specialized individuals perform niche tasks (***Wilson (1980)***; ***Wheeler (1991)***; w***Powell and Franks (2005)***; ***Schofield et al. (2011)***; but see ***Dornhaus (2008)***). Although fixed allocation strategies may be efficient in stable environments, a more flexible approach is needed to respond to changing conditions (***Gordon, 2014, 2016***). Responsive, decentralized changes in task allocation can arise, for example, from individuals with different response thresholds for task-specific stimuli (***Bonabeau et al., 1997***), individuals selecting tasks based on current need or availability (***Tofts, 1993***), state-dependent probabilities to switch or remain in a current task (***Gordon, 1999; Goldsby et al., 2012***), age- or developmentally-related task engagement (***Seeley, 1982; Robinson et al., 1989; Hölldobler and Wilson, 1990***), or a combination of these mechanisms (***Johnson, 2010***). Non-specialized tasks may be distributed widely among individuals, while tasks requiring certain physiological abilities are restricted to individuals able to perform them (***Johnson, 2003; Robinson et al., 2009b***). Across different species of social insects, how and when tasks are allocated among individuals represents a balance between robustness and flexibility in colony function (***Charbonneau and Dornhaus, 2015***).

Task allocation is one means that social insect colonies use to capitalize on favorable conditions to gather sufficient food to survive. Since resource availability can be stochastic, colonies must be responsive to sudden changes in the environment. This includes how to respond, as well as who should respond. In rock ants *(Temnothorax albipennis)*, when additional foragers are needed, workers with the lowest fat content are recruited to foraging (***Robinson et al., 2009b, 2012***). In desert harvester ants *(Pogonomyrmex barbatus)*, interactions activate foragers to make susequent foraging trips (***Davidson et al., 2016; Pagliara et al., 2018***). Honey bees are famed for their ability to stockpile nectar from the environment and convert it to honey; when plants produce copious nectar, a honey bee colony can exploit the favorable foraging conditions by recruiting additional foragers, by having existing foragers perform more trips, or by a combination of the two (***Seeley, 1995***).

In colonies of the honey bee *Apis mellifera* individuals perform different tasks according to multiple factors, including developmental state, genetics, and behavioral feedback mediated by social interactions (***Robinson, 2002; Huang et al., 1994; Beshers and Fewell, 2001; Johnson, 2008b; Wild et al., 2021***). This gives rise to a general tendency for young bees to care for brood in the center of the nest, middle-age bees to perform various tasks throughout the nest, and old bees to forage outside and advertise food sites with waggle dances on the dance floor (***Seeley, 1982***). However, an age-based categorization does not account for how individuals vary throughout their lives, and how previous social and/or environmental experiences may influence task allocation ***Jeanson and Weidenmüller, 2014; Jeanne, 2016***). Moreover, task- or location-based descriptions do not take into account individual movement activity (e.g. speed and spatial localization), which may differ even for individuals involved in the same task (***Crall et al., 2018***).

Several studies have demonstrated that *genetic diversity* enhances colony function (e.g. disease resistance (***Seeley and Tarpy, 2007***) and productivity (***Mattila and Seeley, 2007; Mattila et al., 2012***)), and that there is a genetic basis for honey bee behavior (***Calderone and Page, 1988; Robinson and Page, 1988; Calderone and Page, 1991; Liang et al., 2012***) which can be linked directly to enhanced colony function (e.g. thermoregulation; ***Jones et al. (2004)***). However, we lack a general understanding of how *behavioral diversity* (which may, of course, be related to genetic diversity), and the corresponding individual differences contribute to the flexibility and robustness of honey bee colonies (***Jeanson and Weidenmüller, 2014***). Indeed, task allocation is known to be flexible, with workers reallocating themselves when necessary, thus highlighting the need to use multiple factors in order to describe the behavior of individual bees (***Seeley, 1989b; Robinson, 1992; Huang and Robinson, 1996; Schulz et al., 1998; Johnson, 2003; Wild et al., 2021***).

Gathering data on the lifetime behavior of a honey bee is difficult, and therefore we have a limited understanding of how individuals change their behavior over time. Martin Lindauer, for example, heroically tracked a single worker bee, manually noting all observed activities over 7.5+ hrs each day, for 24 days (***Lindauer, 1952***). This type of laborious individual tracking has formed the foundation of our current understanding of task allocation, whereby individuals perform “task-repertoires” — groups of tasks that are similar behaviorally and/or spatially (***Seeley, 1982; Johnson, 2010***). Fortunately, recent advances in computer vision and machine learning have made automated tracking of social insects feasible (***Mersch et al., 2013; Crall et al., 2015; Wario et al., 2015; Wild et al., 2018; Crall et al., 2018; Gernat et al., 2018; Jones et al., 2020; Richardson et al., 2021; Bozek et al., 2021***). Whereas previous studies rely on ethograms to assign behavior to individuals (e.g. (***Lindauer, 1952; Seeley, 1982; Seeley and Kolmes, 1991; Johnson, 2003; Siegel et al., 2013; Smith et al., 2017; Perez and Johnson, 2019***)), automated tracking minimizes the potential for human-biases. Automated methods also make it possible to track thousands of individuals simultaneously, thus moving from general trends to detailed, long-term, quantification of behavior. Using these techniques, we can both characterise behavioral diversity, as well as investigate the causes and consequences of individual variability and inter-individual differences.

In this study, we analyze the movement of 4,200+ honey bees across 16 age-matched cohorts within a colony throughout an entire summer (July-October 2018). Quantifying the movement of individual bees, as well as spatial maps of the nest environment (i.e. honey stores, brood rearing area, etc.), we identify changes in behavior throughout a bee’s entire lifetime (ca. 15-50 days). We define behavioral metrics related to substrate use and movement characteristics in the nest, use dimensionality reduction and hierarchical clustering to quantify the range of observed behavior, and use these methods to compare the lives of individual bees. We find considerable differences in nest use and movement characteristics among age-matched individuals on any given day. Extending this analysis to the entire lives of individuals, we find that bees not only vary in how early they start to forage, but also have consistent lifetime differences in movement activity and spatial localization. Finally, we use our analysis framework to establish how colony activity shifts during a sudden change in a key environmental characteristic - the availability of food. A massive increase in nectar intake, which tripled the honey storage area in the nest in less than a week, was accomplished with only a small overall change in the distribution of activity across the colony. This demonstrates that distributed changes in the behavior of many bees throughout the nest can facilitate a dramatic colony-level response, while imposing minimal interruption to the allocation of work to other necessary activities.

## Results

### Long-term tracking of individually-marked bees

To obtain the trajectories of individual bees across their entire lifetimes, we tagged and tracked over 4,200 individuals using the BeesBook tracking system (***Boenisch et al., 2018***) (Figure 1A). Newborns were introduced to the 3-frame observation hive every 4-6 days, in cohorts of 200-600 bees. The colony was recorded continuously at 3 frames per second for 86 days, from 16 July to 9 Oct 2018. Each time a new cohort was introduced, we traced the comb contents in the observation hive (as in (***Smith et al., 2016***)) to map the honey stores, brood nest, and dance floor, the latter of which is where foragers advertise food sites. These content maps allow us to determine the spatio-temporal patterns of activity exhibited by bees throughout their lives, in the context of their changing social and structural nest environment (Figures 1B and S1).

**Figure 1.**
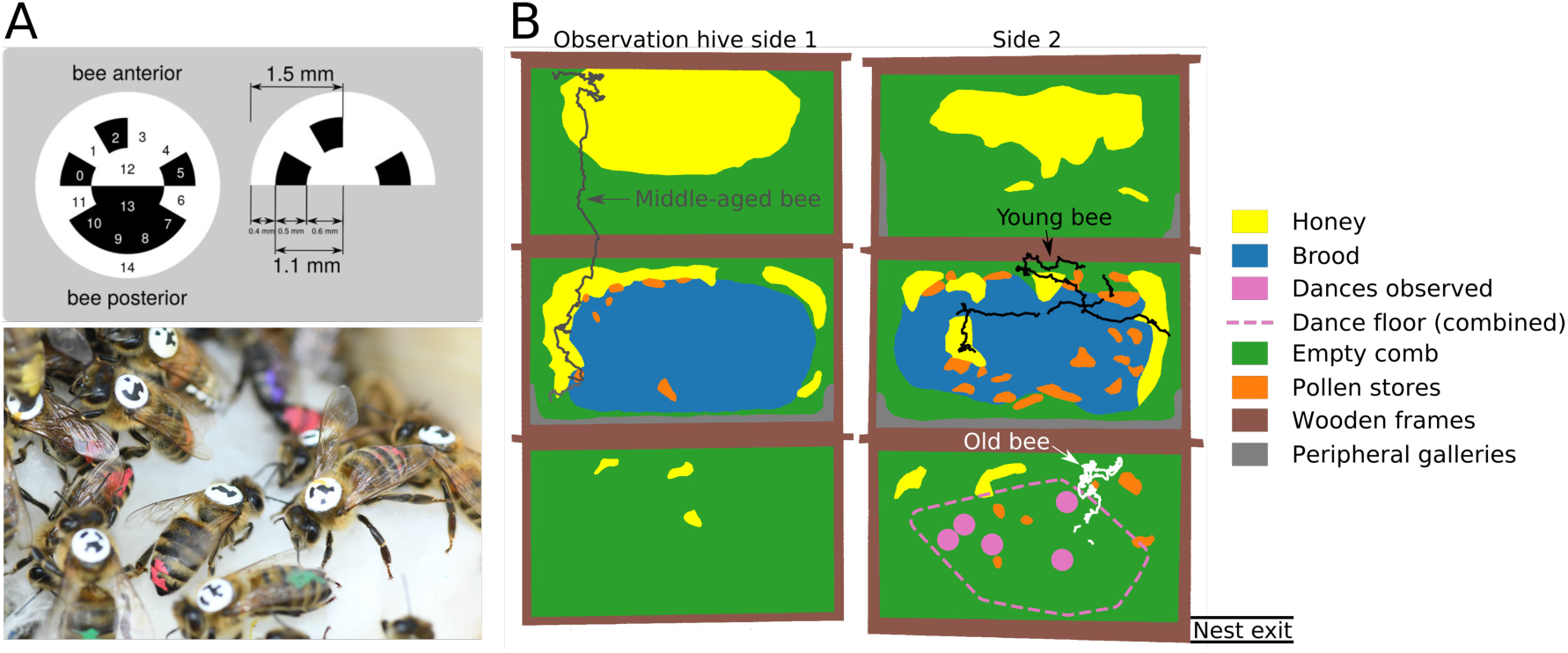
Long-term tracking of bees. (A) Bees were individually marked with barcodes, and tracked using the BeesBook tracking system (***Boenisch et al., 2018***). (B) An example map of the observation hive, with colors to denote different nest substrates. Dots overlaid on the map show trajectories of three representative bees over 2 hours on 17 July 2018: Young bee, age 1 day, in black; Middle aged bee, age 15 days, in gray; Old bee, age 31 days, in white. Nest exit/entrance at the lower right corner.

At any given time, bees on honey storage and brood areas tend to be younger than bees on the dance floor (Figure 2A). As individuals age, they spend more time on the dance floor, and their median speed increases (Figure 2B). These overall trends in nest use match with the well-established sequence of young workers performing within-nest tasks, and old workers foraging outside (***Seeley, 1982; Robinson, 1992***). However, we observe considerable differences among bees within the same age-matched cohort on any given day (Figures 2B, S2). For example, even though both dance floor usage and median speed are positively correlated with age (correlation values of 0.42 and 0.36, respectively; see Figure S3), the correlation between these behavioral metrics is only 0.21 (Figure S3), and is lower still within a certain age grouping (Figure 2C). This highlights the need for more general methods to characterize the differences in observed behavior.

**Figure 2.**
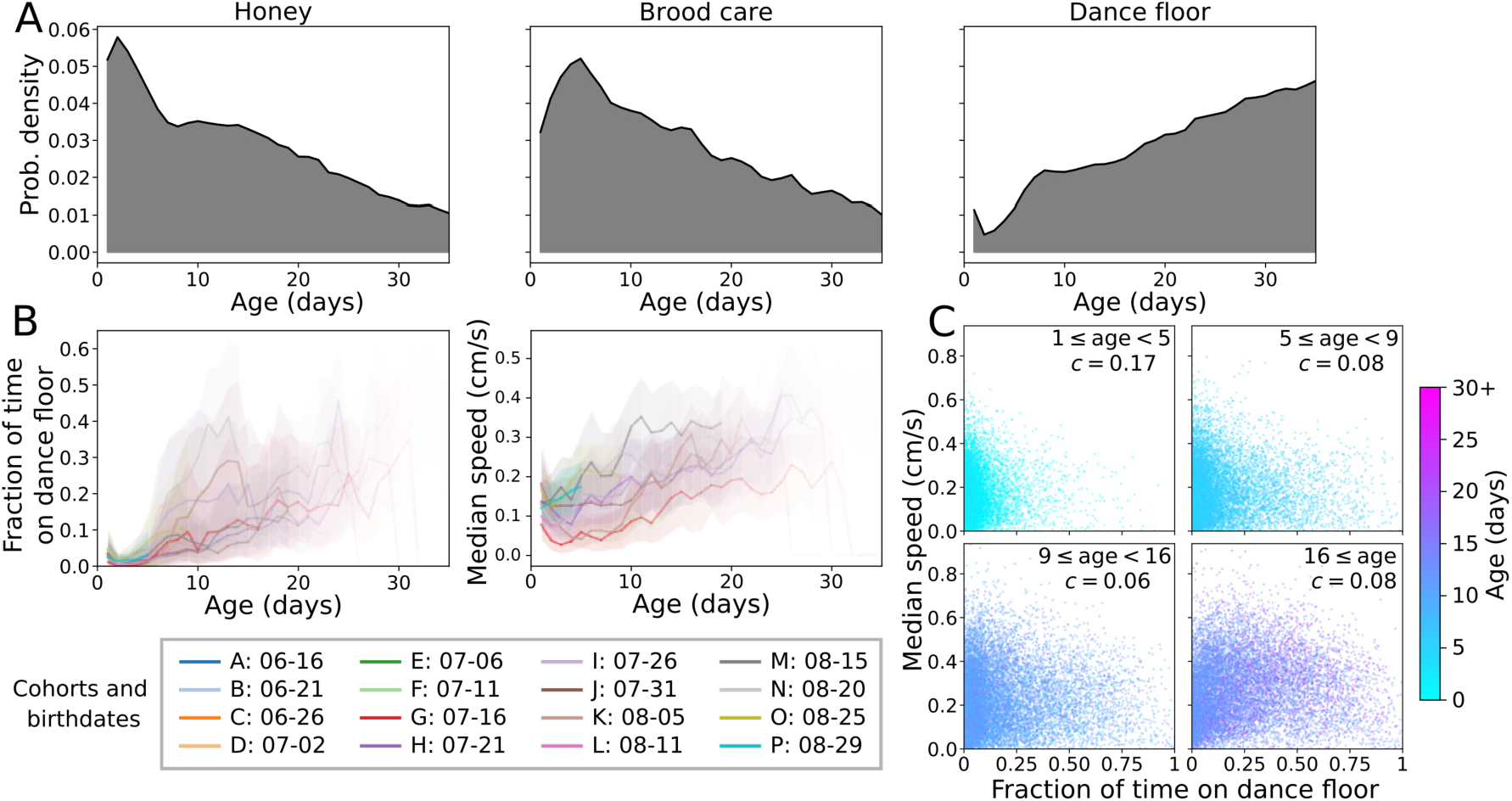
Bee nest usage histograms and changes with age. See also Figures S2, S3. (A) Substrate usage histograms with respect to age. (B) Cohort distributions of substrate usage with age. Colors represent the different cohorts, ordered chronologically by birthday, with corresponding alphabetical names. Lines show the mean and the shaded area shows the standard deviation across bees for each cohort. The transparency is proportional to the fraction of bees in a cohort that lived to a certain age. (C) Plots of the fraction of time a bee spends on the dance floor in a single day versus median speed for that day, grouped by different ages. Points are colored according to the age of the bee on that day. The Pearson (ranked order) correlation *c* between dance floor usage and median speed for bees of that age range is shown on each plot.

### Identifying the structure of individual variation

To quantify the activity of individual bees on a given day, we compute nest substrate usage and movement metrics for each day of the experiment. After filtering to remove dead or unobserved bees (see Methods), the data include 4,201 bees tracked over a focus observation period of 50 days, during which time new cohorts were regularly introduced every 4-6 days. With this, the data includes a total of 53,608 bee-days, where the term “bee-day” refers to the behavior of a single bee on a single day. Quantitatively, a bee-day refers to the behavioral metrics calculated over the course of a day for a single tracked bee in the dataset. After normalizing the behavioral metrics, we use dimensionality reduction to extract the dominant axes of variation and clustering to describe the space of variation. Note that this does not assign specific activities (e.g. cleaning cell) to individuals over time, like an ethogram, but instead uses trajectories and maps of the nest environment to extract salient behavioral features of individual bees each day. Considering all bees on all tracked days, we first describe the metrics associated with different areas of behavioral space. Following this, we use the results to examine variation among age-matched bees, as well as to ask how individual behavior changes over time.

Using principal component analysis (PCA) we first extract the dominant axes of variation. The first PCA component predominantly represents differences in substrate use. The second component is most strongly represented by movement metrics, including speed and dispersion, which is the root-mean square displacement about the centroid of a bee’s trajectory — this is a measure of how wide-ranging each bee is, regardless of where it is located in the nest (Figure 3A). Low dispersion represents staying localized in a single area (i.e. having a small spatial fidelity region, regardless of where that region is located in the nest), while high dispersion represents the tendency to move across multiple areas of the nest (i.e. having a large home range). The first 2 PCA components, which together explain 51.1% of the total variance, show that the main axes of behavioral variation for individual bee-days can be described as foraging versus nest work (PCA 1), and the overall level of movement activity (PCA 2).

**Figure 3.**
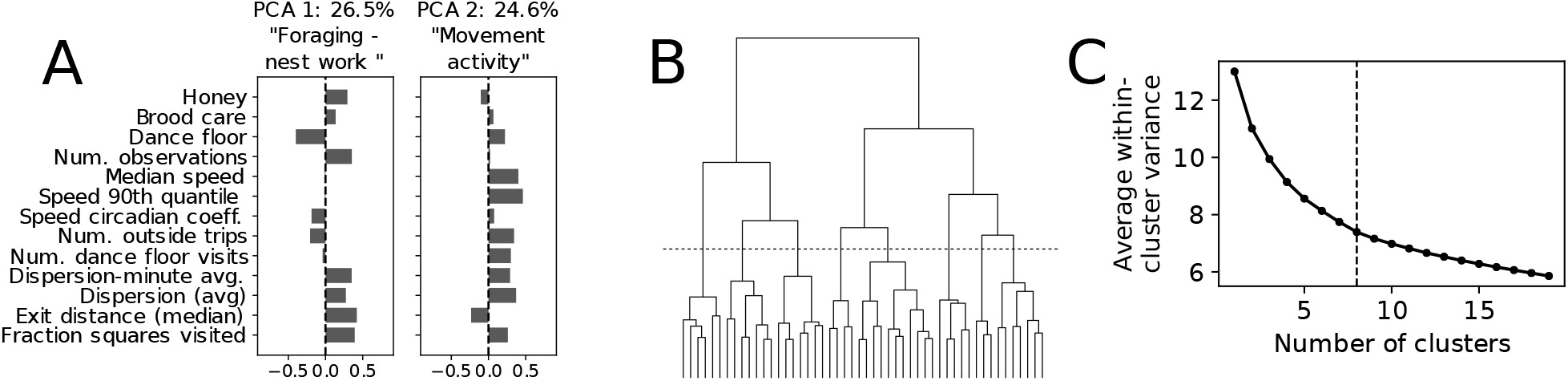
PCA and clustering of individual bee behavior on a given day. See also Figure S4. (A) The first two components from the PCA decomposition of individual bee behavioral metrics on a given day represent the dominant axes of behavioral variation (see Figure S4 for all components). (B) Clustering dendogram and (C) average within-cluster variance as a function of the number of clusters (bottom). In both, the dashed line shows the distance cut-off for the 8-cluster division used in subsequent figures.

We next apply Ward hierarchical clustering (***Ward Jr, 1963***) to group bee-days with similar behavioral metrics, and to ask if distinct behavioral clusters exist. The dendogram structure and the continuous decrease of the within-cluster variance with the number of clusters suggest there are not distinct divisions between clusters, but rather that the overall variation in the data has a continuous structure. Because of this, we use clustering as a tool to describe differences in behavior, not to suggest the existence of distinct behavioral clusters. While the number of clusters in a continuous data space is arbitrary, fewer clusters can be used to describe coarser trends, while more clusters reveal finer structure in the data. Based on the dendrogram structure, we focus on 8 clusters as a useful division in the data (Figure 3C);an expanded feature diagram with different numbers of clusters is shown in Figure S11.

To visualize the full range of variation in the data, we use t-distributed stochastic neighbor embedding (t-SNE; ***Maaten and Hinton (2008)***), which assigns each datapoint a location within a two-dimensional space. Because we use the first two PCA components to initialize the t-SNE, the global structure of the t-SNE embedding reflects these two dominant axes of variation (Figure S5). However, since t-SNE compresses the high-dimensional space into only two dimensions, the t-SNE embedding positions reflect more than just the first two PCA components. We use this visualization to describe observed differences in bee-days (Figure 4).

**Figure 4.**
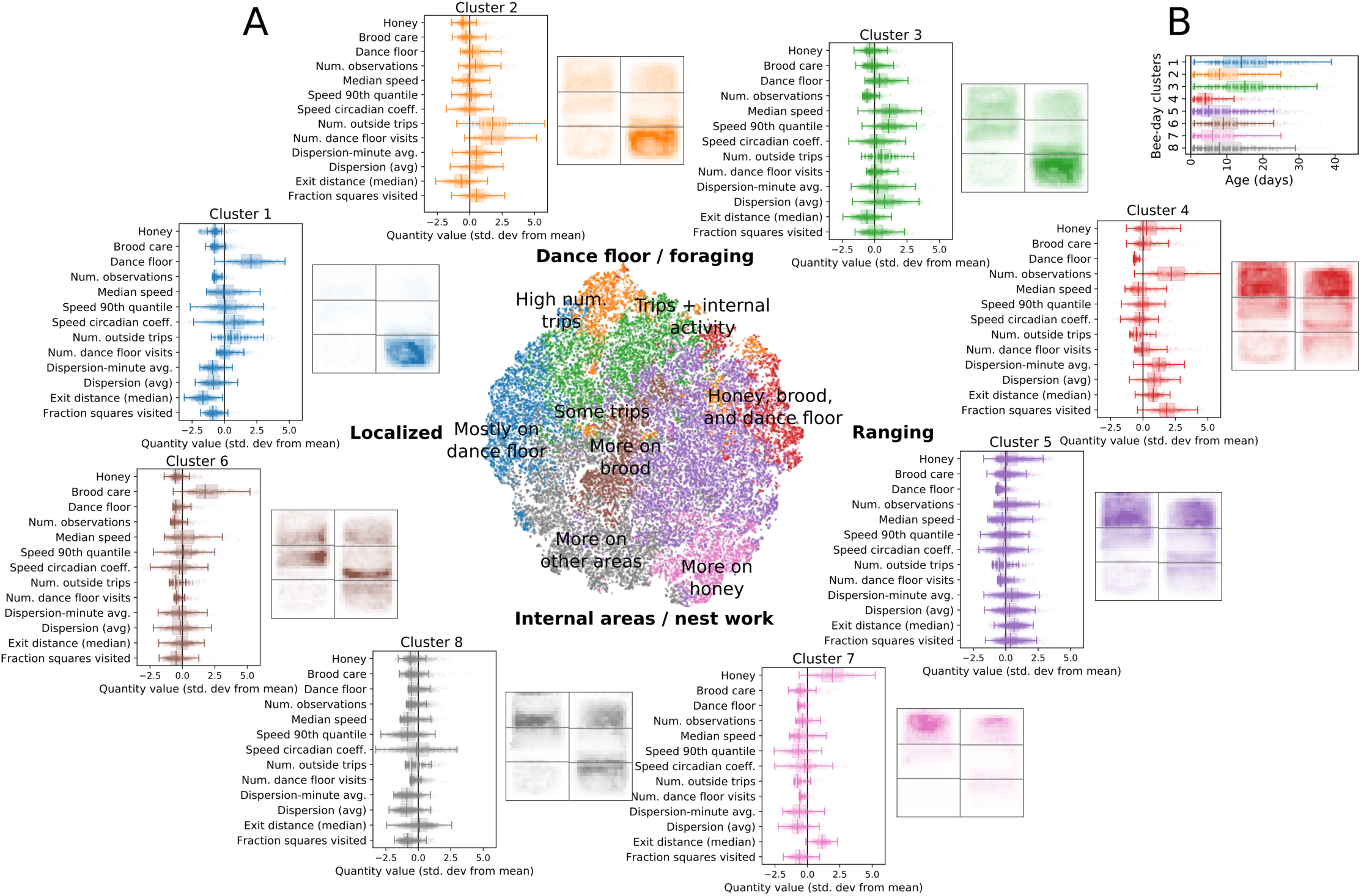
Differences in observed bee-day behavior. See also Figure S11. (A) Distributions of bee-day data using 8 clusters, showing behavioral metrics and average nest location histograms. Colors highlight the different groups on the t-SNE embedding (center). Nest histograms reflect the 6-frame layout of the observation hive shown in Figure 1B. (B) Age distributions of each bee-day cluster.

“Foraging” bee-days are characterized by more time spent on the dance floor, more outside trips, and more dance floor visits;”Nest work” bee-days are characterized by more time spent on brood or honey areas. Both foraging and nest work activity vary continuously along the axis of low to high dispersion;for example, some foragers exhibit lower dispersion and are observed mostly on the dance floor (cluster 1 - blue), whereas others exhibit higher dispersion and are observed in other nest areas (clusters 2 and 3 - orange, green). Nest workers may also exhibit low dispersion (cluster 8 - gray), intermediate dispersion (cluster 5 & 6 - purple & brown), or high dispersion (cluster 4 - red), along a continuum (Figures 4A, S5). Although speed and dispersion are correlated (Figure S3), and are weighted similarly in the second PCA component (Figure 3A), the nonlinear embedding highlights instances where speed and dispersion differ (c.f. cluster 3 and cluster 4 in Figure 4A).

Some bees with low dispersion spend time on areas of the nest other than honey, brood, or the dance floor (cluster 8 - gray). Note that substrate use is normalized across all nest areas shown in Figure 1, and is mutually exclusive, such that if a detected bee is not on honey, brood, or the dance floor, then she is on “other” nest areas (see Figures 1, S1, and Methods). In addition to the two primary axes of variation (foraging-nest work; low-high dispersion), we also observe bee-days that are distinguished by a high number of outside trips and dance floor visits (cluster 2 - orange), more time in the brood area (cluster 6 - brown), and more time in the honey store areas (cluster 7 - pink). Although the age distributions for each cluster overlap (Figure 4B), the median ages in clusters 1 and 3 are the highest (foraging bee-days), while the median ages of bees in clusters 4 and 5 are the lowest (nest work bee-days with high dispersion). Overall, these results show how differences in bee behavior are described by a continuous distribution along dominant axes that correspond to nest substrate use (e.g. time spent on the dance floor) and movement metrics (e.g. dispersion throughout the nest).

### Age-matched differences and changes over time

Bees change their behavior over time due to internal processes such as physiological development, as well as due to interactions with other bees, and environmental factors (***Robinson, 1992; Amdam and Omholt, 2003; Johnson, 2003, 2010***). To investigate how bees change their activity patterns with age and over time, we compare age-matched individuals using the embedding and clustering results, which aggregate all individual quantities into a single embedding. Younger bees tend to exhibit behavior associated with nest work (clusters 4-8: red, purple, brown, pink, gray), while older bees tend to exhibit behavior associated with foraging (clusters 1-3: blue, orange, green) (Figure 5A).

**Figure 5.**
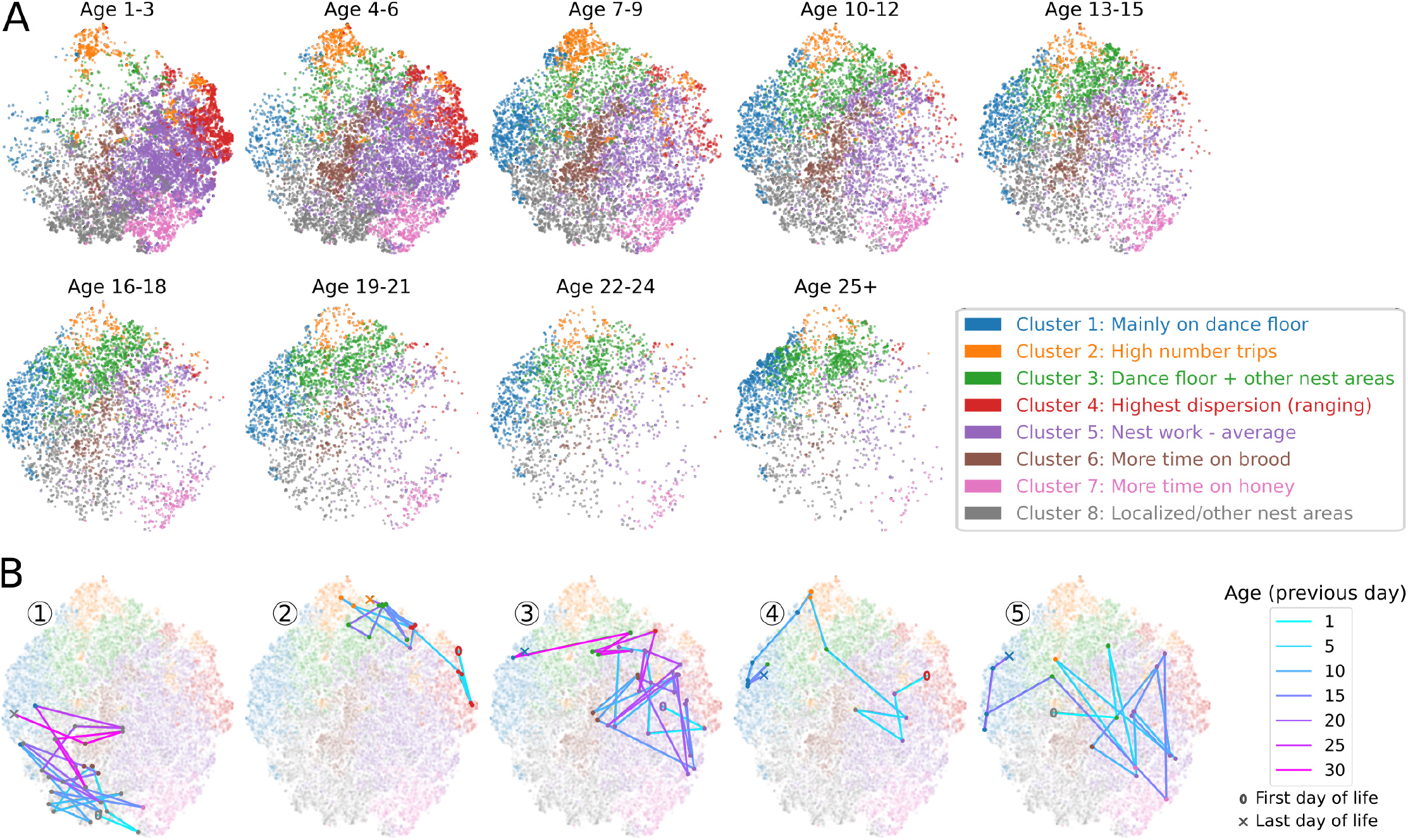
Changes in activity with age and over time. The colors in all panels correspond to the behavioral clusters shown in Figure 4. (A) t-SNE embedding points for bees of a particular age. (B) Example behavioral trajectories of individuals bees in t-SNE space. In each, the background shows the t-SNE space, the colored points show the behavioral category of the bee on a certain day, and the lines connect subsequent days, with shading based on the age of the bee on the previous day. The numbers 1-5 identify the individual bees marked in Figure 6.

We can also examine how individual bees change their behavior with age, by plotting the behavioral embedding for each day of their lives. This shows that individuals take different paths through behavioral space (Figure 5B). Moreover, even though bees change behavior with age, some bees have consistent differences in behavioral metrics that persist throughout their entire lives. For example, bees 1 and 2 in Figure 5B occupy different regions of behavioral space, with bee 1 generally having low dispersion, regardless of its age, and bee 2 having high dispersion, regardless of age. Conversely, bees may occupy similar regions of behavioral space, but may change behavior at different rates. This is true for bees 3 and 4 in Figure 5B, whose lives occupy similar regions in behavioral space, but differ in the rates that they transition through this space. These examples, along with observations that behavior on subsequent days tends to be similar to the previous day (Figure S6), demonstrate that there are patterns in how individual bees change their behavior as they age.

### Behavioral clustering of bee-lives

To quantitatively compare how different bees change behavior as they age, we use a procedure similar to that used for single days. However, now instead of a single day, each point represents the entire life of an individual bee — we refer to this as a “bee-life”. Quantitatively, a bee-life is defined by the behavioral metrics for each day of the bee’s life. Individual bees have different lifespans; to compare bee-lives we include bees that have data for at least 10 days, and use per-age average values of each metric to fill in values of the bee-life matrix for the purposes of PCA and clustering, for days where a bee was not observed or where they had already died (see Methods). Applying PCA to the bee-life data, the first mode predominantly shows consistent differences in movement activity, including speed and dispersion, that persist across the entire life of a bee. This differentiates bees that, regardless of age, tend to have high dispersion (e.g. Bee 2 in Figure 5B) or low dispersion (e.g. Bee 1 in Figure 5B). The second mode reflects differences with respect to how behavior changes with age, highlighting differences between bees that begin foraging-related behavior at an early age (e.g. Bee 4 in Figure 5B), versus those that live longer and begin foraging at a later age (e.g. Bee 3 in Figure 5B). This mode represents early-to-forage behavior, with an increase in outside trips, dance floor visits, and dance floor usage that begins at an early age (Figure 6A).

**Figure 6.**
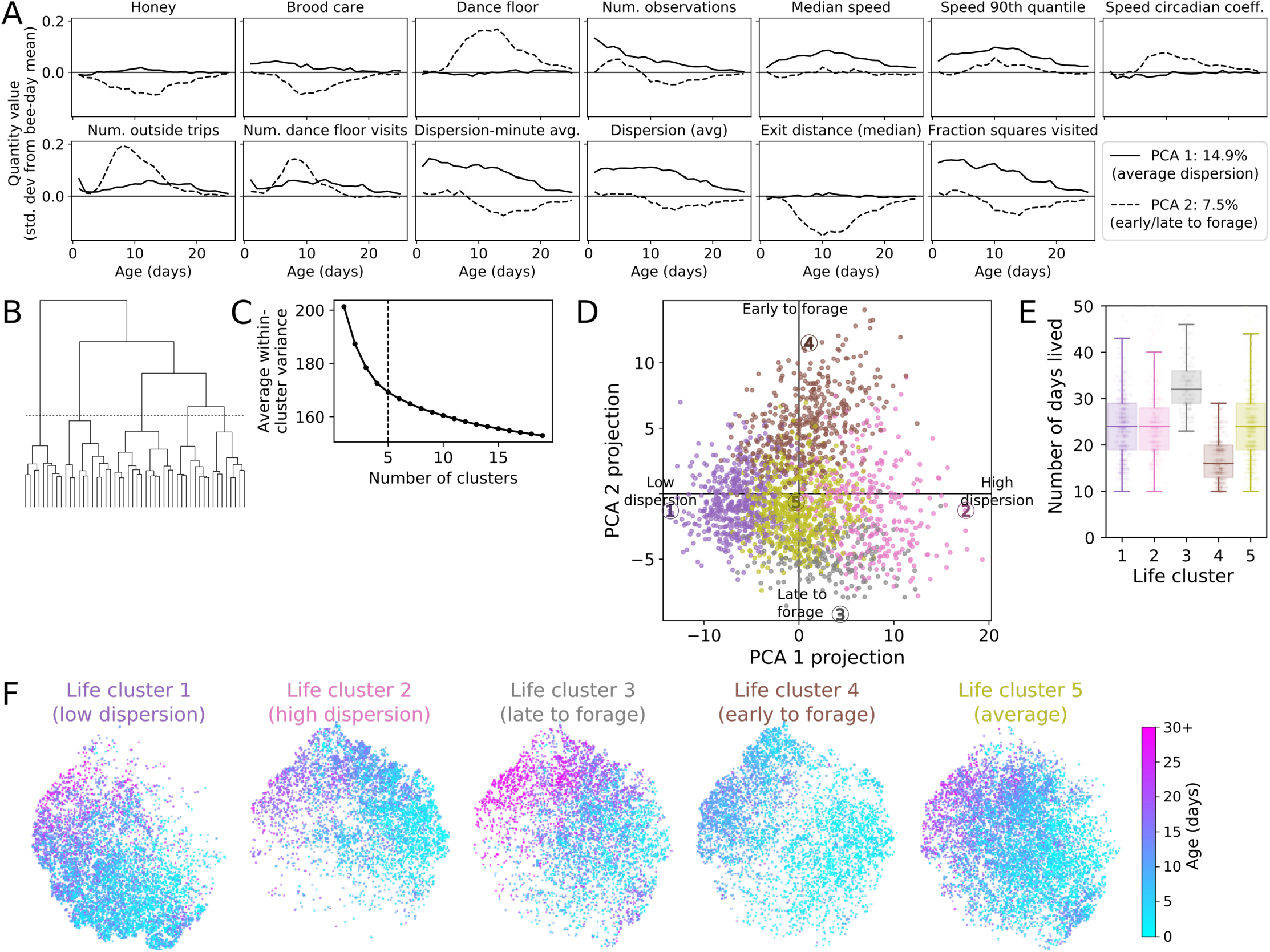
Differences in behavior over a bee’s life. See also Figures S7, S9, S10, (A) PCA decomposition of bee lives shows the dominant modes for how the behavioral metrics change over time. Plots show each PCA mode plotted in terms of behavioral metrics with age, using normalized quantities with the same units as Figure 4 (i.e. zero represents the mean of a certain behavioral metric across all bee-days). (B) Clustering dendogram and (C) average within-cluster variance as a function of the number of clusters (bottom). In both, the dashed line shows the distance cut-off for the 5-cluster division used in this figure. See Figures S9, S10 for results with an 8-cluster division. (D) A plot of individual bee-lives projected onto the first two PCA modes. Each point represents the life of a single bee. The colors correspond to a 5-cluster division, identified via Ward hierarchical clustering. The labels 1-5 identify the individual bees shown as examples in Figure 5. (E) Distributions of number of days lived for bees in each life cluster. Note that bees are only included if they lived at least 10 days. (F) t-SNE embeddings in the bee-day space (see Figures 4 and 5), selecting points associated with bees in each life cluster. Points are colored by age.

We use clustering on the bee-life data, which includes the behavioral metrics for each day of a bee’s life (see Methods), in order to group together individuals with similar lives, and describe differences in the overall space of lifetime variation. The average within-cluster variance continues to decrease with the number of clusters, and individual bees are distributed continuously along each of the PCA axes, thus suggesting that distinct clusters do not exist. We use a division into 5 clusters in order to describe lifetime differences along the dominant two PCA modes (Figure 6B-C); because the first two PCA modes represent the largest fraction of the variance, further divisions into more clusters still mainly reflect differences along these axes (Figures S9, S10). We use the PCA components as shown in Figure 6A to examine how the behavioral metrics during the life of bees in each life cluster project onto these modes. Life clusters 1, 2, and 5 predominantly represent differences in the projection onto PCA mode 1. Life cluster 1 bees have lower average dispersion and speed values, life cluster 5 bees have intermediate values, and life cluster 2 bees have higher average values of dispersion and speed (see also Figure S9). Figure 6F uses the bee-day t-SNE embedding to compare the behavioral space occupied by bees in each life cluster, with points colored by age. This shows in particular that differences between bees in life clusters 1 and 2 are reflected in the behavioral space of the bee-days that make up the lives of individuals in each cluster (Figure 6F).

Using the PCA modes shown in Figure 6A, bees in life cluster 4 have positive projections onto PCA 2, which is the “early-to-forage” mode; these bees also lived shorter lives than other bees (Figure 6D-E). Although we did not gather direct observations of foraging trips for individual bees, the higher numbers of outside trips, dance floor visits, time on the dance floor, and the shorter life span are characteristic of forager bees (***Visscher and Dukas, 1997***), and thus we refer to bees in life cluster 4 as “early-to-forage”. Precocious foraging, which is similar to the “early-to-forage” individuals that we observe, can be induced via hormone treatments (***Robinson et al., 1989***), infection (***Woyciechowski and Moroń, 2009***), or colony demography (***Huang and Robinson, 1996***), but here we see that such individuals exist even in unmanipulated colony conditions, similar to recent findings in ***Wild et al. (2021)***.

Conversely to life cluster 4, bees in life cluster 3 have a negative projection onto PCA mode 2, and tend to live longer than other bees. Comparing life clusters 3 and 4, the main difference is not in the overall region of behavioral space that is used, but rather in how quickly bees transition through this space (Figure 6F). For both life clusters, bees transition to foraging behavior as they age (note that foraging bee-days are located in the upper-left area of the bee-day behavioral space; c.f. Figure 4). Figure 6F thus shows that while bees in life cluster 1 are “early-to-forage”, bees in life cluster 4, i.e. bees with negative projections onto PCA 2, can be described as “late-to-forage”.

The differences described by life-PCA component 2 agree with recent work which found differences among individual bees in developmental trajectories, as described in the age at which they change interactions patterns and nest substrate usage (***Wild et al., 2021***). Moreover, ***Wild et al. (2021)*** found that developmental trajectories began to significantly diverge at approximately six-days of age. We also find that “early-to-forage” bees tend to show differences in behavior near this age (Figures 6A, S10), thus suggesting a potential critical point in the early life development. Other work with bumblebees has found consistent lifetime differences in movement activity among individuals (***Crall et al., 2018***). Here, we also see this with honeybees, as described by life-PCA component 1. Overall, we see that individual bees differ in the behavior they exhibit over the course of their lives, and that the dominant axes of variation are: (1) individual differences in movement metrics (PCA 1 - average dispersion values), and (2) how early a bee starts to forage (PCA 2 - early or late to forage).

### Colony activity during a nectar flow

Honey bees must capitalize on favorable conditions - such as when new flowers come into bloom - in order to gather sufficient food to survive. Between 26 and 31 August 2018 (experimental days 41-46), we observed a dramatic increase in nectar availability (a “nectar flow”), wherein the colony more than tripled the size of its honey stores (436 to 1429 cm^2^) and increased its weight by 1.25kg (Figure 7A). Given that a forager carries 55-65mg per trip (***Seeley, 1994***), this represents at least 3000 additional successful foraging trips per day. Considering all of the tracked bees in the colony, we see that the total number of outside trips and dance floor visits increase by several thousand when the nectar flow period begins (Figure 7B).

**Figure 7.**
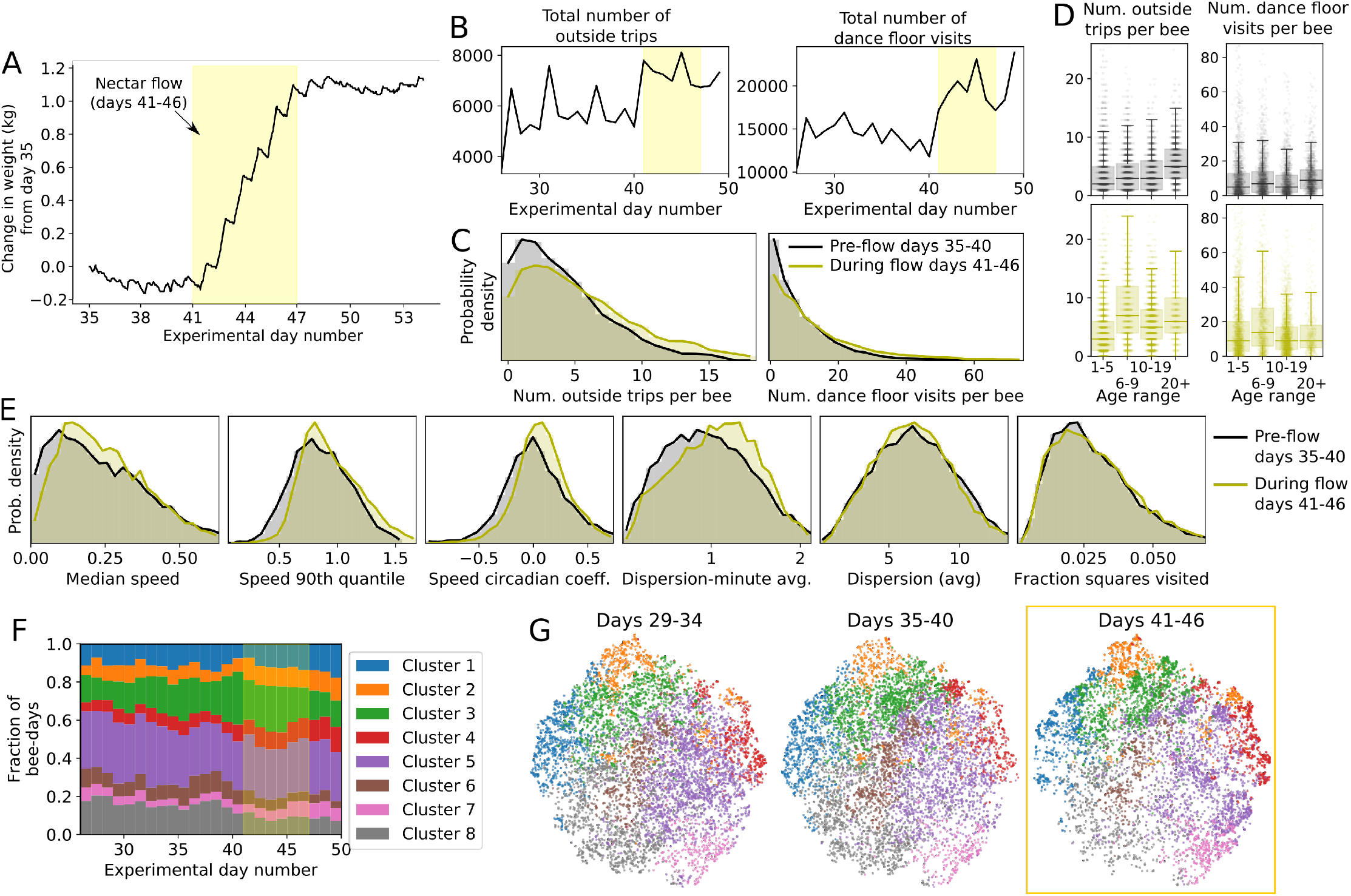
Nectar flow and changes in bee activity. (A) Colony weight change from day 35, with the nectar flow shaded in yellow. (B) The total number of outside trips and dance floor visits per day for all tracked bees in the colony during the second half of the focus observation period. The nectar flow is shaded in yellow. (C) The distributions of number of outside trips and dance floor visits for individual bees on days before the nectar flow versus during the nectar flow. (D) The same data as (C), but shown as boxplots grouped by the specified age ranges of bees alive during each corresponding time period. (E) Distributions of movement metrics before and during the nectar flow. (F) Fraction of bee-days in each cluster over time (colors match clusters in Figure 4). The shaded yellow area indicates the nectar flow. (G) t-SNE embedding of the behavioral data before and during the nectar flow (yellow box), shown in consecutive 6-day periods. For reference, the labels at the leftmost image show the dominant axes of behavioral variation. Note that because the age distribution of marked/tracked bees is skewed towards younger bees in the first half of the recording period, we focus on comparing colony activity from experimental day 26 (11 August 2018) onward.

How is this change distributed among individuals? The increase in outside trips and dance floor visits is accomplished by a shift in the distributions of activity per individual bee, which represents a distributed re-allocation of activity (Figure 7C). Although we did not make direct observations of nectar gathering- and processing-related activities, which include foraging trips, waggle dances, and nectar offloading events, the number of outside trips and dance floor visits serve as reasonable proxies for these activites (***Seeley, 1995; Nieh, 1998; Wray et al., 2008***). Given these proxies, the distributed shift in Figure 7C represents both (1) more bees being involved and (2) involved bees having higher activity compared to before the nectar flow. Dividing the distribution into bees of different age groups, we see that although the median number of outside trips and dance floor visits increased for bees of all ages during the nectar flow, the largest change, when comparing pre-flow to during-flow distributions, is seen among bees ages 6-9 (Figure 7D).

We also found that while daily dispersion and fraction of the nest visited did not change during the nectar flow, bees did, on average, increase their speed and their short-timescale, minute-averaged, dispersion (i.e. an increase in short burst activities; Figure 7E). We did not observe a large shift in the cluster assignments for the daily work activity of individuals during the nectar flow (Figure 7F,G). Together, this demonstrates that the colony is able to accommodate the increased foraging activity by distributing the workload among many bees, presumably to maintain other tasks and activity in the nest when doing so.

## Conclusions

A honey bee colony represents a special level of biological organization — a “superorganism”, where individual workers form a cooperative group to propagate their genes (***Smith and Szathmary, 1995; Hölldobler et al., 2009***). Individual task allocation enables the colony to be flexible in response to changing conditions, yet robust to the maintenance of other colony functions. Using individual tracking data from 4,200+ bees, we show how honey bees differ in behavior at the timescale of both days (Figure 4) and lifetimes (Figure 6). Across both time-scales, the dominant differences among individuals are in nest use (in particular, time spent on dance floor), and movement patterns (speed and dispersion, which determine how much of the nest an individual samples). Beyond these dominant axes of variation, we used dimensionality reduction and clustering to describe further differences due to number of outside trips, number of dance floor visits, time on honey, brood, or other areas, and combinations of these (Figure 4). Examining entire lifetimes, we see that some individuals show consistent differences in movement characteristics, or begin to forage at different ages (Figure 6). Overall, our long term tracking results present a detailed picture of how individuals differ in behavior from day-to-day, over entire lifetimes, and how this behavior is organized at the colony level.

While previous work has used ethograms to define categorical age-based labels such as nurses, middle-aged bees, and foragers (***Lindauer, 1952; Seeley, 1982; Seeley and Kolmes, 1991; Johnson, 2008a***,b, ***2010***), our analysis shows that these categories are a partial description, representing slices within a continuous behavioral space (Figure 4). An important goal for future work will be to compare our results to approaches that use ethograms to manually assign behaviors and task repertoires. A further open question pertains to how individual variation may be adaptive in certain environmental conditions, developmental states, or group functions ***Jeanson and Weidenmüller, 2014***). The question of optimal inter-individual differences and variability in response also applies to bio-inspired robotics, where current work seeks to incorporate these mechanisms to achieve flexible and adaptive functionality (***Dorigo et al., 2013***). Our results show the structure of behavioral variation of individuals in a honey bee colony (Figure 4), and suggest that continuous variation may enable a distributed response such that the colony can maintain vital functions while adapting to changing conditions (Figure 7).

In the analysis of the activity patterns of bees over their entire lives, we see that some individuals are *consistently* different: Figure 6 shows that some bees have consistently higher (or lower) dispersion over their life (Figure 6). While previous work has found consistent differences in movement activity of bumblebees from day to day (***Crall et al., 2018***), to our knowledge, this is the first description of a honey bee movement metric that shows lifelong consistency.

Social insect colonies show flexibility to adapt behavior to changing conditions, yet also robustness to maintain important colony functions. We studied how a honey bee colony responds to changing environmental conditions by examining individual bee behavior during a naturally occurring dramatic change in nectar availability, which resulted in the colony tripling the size of its honey stores. During this period, we see that the increase in nectar coming into the colony is accommodated by a distributed task load, whereby many individuals exhibit small increases in the number of outside trips and the number of dance floor visits (Figure 7). Although bees of all ages increased their activity with respect to these metrics, the change was largest for bees ages 6-9 (Figure 7D). Bees of this age range during the nectar flow mostly belong to cohort N; while we see that bees in this cohort tend to show more “early-to-forage” behavior than bees in other cohorts (Figure S8A), further experiments are needed to ask whether such a difference is driven by genetic factors (e.g. source colony differences - Figure S8B) or environmental factors. The observed distributed increase in activity during the nectar flow demonstrates how a colony can adapt to changing conditions without a large-scale reorganization — small changes, when compounded, are sufficient. While collecting nectar is essential for the long-term survival of the colony (***Smith et al., 2016***), a distributed system may also prevent the colony from risking over-investment that would jeopardize other colony requirements, thus balancing a trade-off between flexibility and robustness. Indeed, a honey bee colony represents a biological entity that must be resilient to multiple co-occurring challenges both within and outside the colony. These evolutionary pressures have led to inter-individual differences and variability in behavior among the individuals that make up a colony.

## Methods

### Observation hive and nest maps

This research was conducted at the University of Konstanz, Germany (47.6894N, 9.1869E). On 10 June 2018, the observation hive was installed with a single queen, 2,000 unmarked workers, and three frames of mixed brood and honey (“Deutsche-Normal” frames: 395 × 225 mm, observation hive: 490 × 742 mm). From 16 July to 29 Aug 2018, every 4-6 days, we individually marked and introduced 200-600 newborn honey bees to the observation hive (total bees tagged: 5343). Although tracking data was obtained continuously until 9 October 2018, we perform our analysis on a focus observation period of 16 July - 3 September, during which new cohorts were regularly introduced. Newborns were hatched overnight in an incubator kept at 34°C and 50 %RH, and marked the following morning with individual BeesBook tags (***Wario et al., 2015; Boenisch et al., 2018***). From 16 July to 3 Sept 2018 (50 days) we recorded the obervation hive at 3 frames per second using four Basler acA4112-20um cameras fitted with Kowa LM25XC lenses. The colony was illuminated with infrared light (850nm 3W LED’s), which is invisible to honey bees (***Peitsch et al., 1992***). The entire recording rig (observation hive, cameras, lighting) was kept in the dark, to mimic the natural conditions of the honey bee nest. Workers had free access to forage outside, through a entrance tunnel (2-cm diameter). To keep track of the colony’s weight, the observation hive was kept on a scale which logged its weight every hour (10g sensitivity, Wolf Waagen GmbH). To create a map of the nest, every 4-6 days we traced the contents of the observation hive onto plastic sheets by outlining the following: honey storage, pollen storage, brood, empty comb, wooden frames, peripheral galleries, and dances observed on the dance floor (as in (***Smith et al., 2016***); Figures 1B, S1). These plastic sheets were then scanned with an architectural scanner (Ruch-Medien, Konstanz), and digitized. By overlaying the bee trajectories upon the maps, we determined what type of nest environment an individual experienced (Figure 1B).

### Data processing and quantities to describe behavior

Using the BeesBook system, the raw image data were processed to detect and decode the individually marked bees (***Boenisch et al., 2018; Wild et al., 2018, 2021***). For each individual, its tag id, id detection confidence, position, and orientation were tracked over time, and stored in a PostgreSQL database. The death date of each marked individual was estimated using a Bayesian changepoint model (as in (***Wild et al., 2021***)). This method accounts for a low rate of erroneous detections in bees that have already died, and time periods when individuals are observed less frequently or not at all (e.g. while foraging). An individual’s death date was used as a cutoff for including data in subsequent calculations.

We processed the trajectory data to obtain the quantities used in the subsequent analyses by first averaging over 1-hour time bins and saving the quantities of interest for each individual bee. All data points used in the analysis were above a detection confidence threshold of 0.8.

We chose metrics that represent use of the nest and space within the nest (Num. observations, time on honey, brood, or dance floor), spatial localization (dispersion - average, dispersion - minute average, fraction of squares visited), and activity levels (median speed, speed 90th quantile, speed circadian coefficient, num. outside trips, num. dance floor visits). Although some of these metrics have similar trends (e.g. dispersion and fraction of squares visited, or speed and speed 90th quantile - see correlations in Figure S3), they nonetheless represent different aspects of behavior, and we use the approach of combining multiple different metrics in order to obtain results that are robust to particular parameter choices. For each hour of each day, we calculated behavioral metrics for each bee that had a minimum of 10 detections in that hour. For ease of processing the large amount of data, all quantities were calculated for each hour of each day. For the metrics of substrate usage, speed, exit distance, and fraction of squares visited, the per-day quantity was then determined as a weighted average across the hours in the day. For number of observations, number of outside trips, and number of dance floor visits, the per-day value is a sum across our hours. For dispersion - minute average, we used a non-weighted average in order to give a different representation compared to the average dispersion.

#### Substrate usage

is calculated using the comb substrate maps shown in Figure S1, grouping together capped and young brood into a single category. Note that dances were observed only within a limited time range (pink circles in Figure S1), but all occurred in a similar area. Defining the dance floor based on only direct observations would be overly restrictive, so we defined the dance floor area using a complex hull that contains all dances over the entire observation period (dashed pink line in Figure S1). Because the comb contents changed over time, and were not measured each day, we calculated substrate usage by linearly interpolating an individual bee’s substrate usage fraction between values calculated using the substrate maps on the measurement days before and after the day in consideration. To illustrate this procedure, consider the day July 18, which has the closest measurement days of July 16 and 21. Denote the comb map from July 16 as *A*, and the comb map on July 21 as *B*. We first use the trajectory coordinates of the bee on July 18 to calculate two different approximate usage fractions: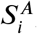, which is the fraction of time spent on substrate *i* as determined using map *A*, and 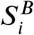, which is the fraction of time spent on substrate *i* as determined using map *B*. The estimated substrate usage fraction for July 18 is calculated is calculated as a weighted average of these values:

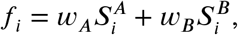

where for this example the weights are *W*_*A*_ *=* 0.6 and *w*_*B*_ *=* 0.4, because the first comb measurement day is closer than the second to July 18. The nest comb contents over time were also determined by this same linear interpolation method between nest content measurement days.

#### Number of observations

is the total number of detections with confidence interval over the 0.8 detection threshold.

#### Median speed

is determined from speed, which was calculated using the distance traveled divided by time interval between detections. We omitted instances where the bee switched sides of the comb, as well as when the time between detections was > 1 second. Similarly, **speed 90th quantile** is calculated using the distribution of speeds.

The **speed circadian coefficient** is calculated as a correlation of median speed over the day with a daily rhythm that follows the sun. We approximate the daily rhythm with a sine curve of sin((*h* − *m*)/(2 *π*), where *h* is the hour of the day, and *m* is chosen so that the maximum of the curve coincides with the highest sun position of the day, which was approximately 13:30 CEST during the observation period. The circadian coefficient is then calculated as

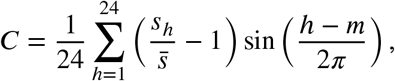

where 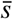 is the hour-averaged speed. With this normalization the coefficient satisfies −1 ≤ *C* ≤ 1.

The **number of outside trips** is estimated by counting instances where a bee was detected in the observation hive frame nearest to the exit, and then subsequently not seen for 10 minutes.

The **number of dance floor visits** is the number of times that a bee entered the dance floor from another substrate.

#### Dispersion - average

is the root mean square distance from the centroid of the x-y coordinates, calculated by neglecting which side of the comb the bee is on, i.e. considering motion in a 2D plane and neglecting the 3rd dimension that describes whether the bee was detected on the front or the back of the observation hive. The average dispersion was calculated by first calculating per-hour dispersion, and then taking a weighted average across the hours of a day. **Dispersion - minute average** was calculated similarly, but using the centroid of the x-y co-ordinates for each minute, and then averaging by equally weighting each minute (instead of taking a weighted average according to number of observations).

The **median exit distance** is determined by calculating the shortest distance to travel to the exit (which is located in the lower right corner), accounting for possible routes to switch sides, but not adding any extra distance for a switch of sides.

The **fraction of squares visited** is determined by first dividing the nest area into discrete spatial regions of 2cm×2cm (the same grid size used in the spatial histograms shown in Figures 4 and S11), and then counting the fraction of bins with at least one observation.

### PCA and clustering on bee-day behavioral metrics

Using the behavioral metrics (honey, brood care, and dance floor substrate usage; number of observations; median speed and speed 90th quantile; speed circadian coefficient; number of outside trips and dance floor visits; dispersion average and minute average; exit distance; fraction of squares visited), we create a data matrix *M*_*ij*_, where each row *i* represents one bee-day, and columns *j =* 1..13 are the different quantities. A bee-day is only included if that bee was alive on the given day and had more than 1,000 detections over the whole day. This represents a total time observed of 5.5 minutes during a day; using this removes 7948 bee-days with few observations (results are qualitatively similar whether these are included or not). In addition, we do not include bees on the first day they were introduced, because on this day there were not observed for a full 24 hours. With this criteria *i =* 1..53608 bee-days are included in the analysis. Although the total number of tagged bees was 5, 343, the bees in cohorts A-F were tagged before filming began, and some died before 16 July. Due to this, and after filtering, we include data from a total of 4, 201 unique bees in the analysis (the number is 4, 322 before filtering for few observations).

Note that the nest contents - in particular the size of the honey and brood areas - change over time (Figure S1). We wish to account for this change in order to focus on variation among the activity of bees in the nest at a given time, instead of changes in substrate usage that result from a different nest composition. For honey and brood areas, we account for this by subtracting the nest content fraction from the individual bee substrate usage fraction for each day. The dance floor is unaffected, since it is defined as the same area over the course of the observation period.

Following standard procedures, we normalized the data matrix **M** so that the column mean is zero and the column standard deviation is 1. We then performed principal component analysis (PCA) on the resulting matrix to obtain the components shown in Figures 3A and S4. The result of PCA is a matrix *U*_*ij*_, where *i* represents bee-days and *j =* 1..13 for the PCA components (corresponding to the total number of behavioral metrics).

Next, we perform Ward hierarchical clustering, implemented in Python in the package `scipy.cluster.hierarchy`, to obtain the clusters shown in Figures 3 -4. We use t-SNE embedding (***Maaten and Hinton, 2008***) implemented in `sklearn.manifold`, with parameters of perplexity=30 and n_iter=1000, with initial conditions set by the first two PCA dimensions to obtain the bee-day embeddings shown in Figures 3, 4, 5 7, S5, and S6.

Ward clustering minimizes the overall within-cluster variance. For *n* clusters, this is calculated as

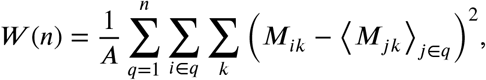

where *A =* 53608 is the number of rows in **M** (i.e. the number of bee-days), and ⟨ · ⟩_*j∈q*_ represents an average over the indices *j* that are elements of cluster *q*. This is shown in Figure 3C.

### Bee-life

The results in Figure 4 treat each day for each bee separately, and each row of **M** represents one bee-day. Building on this notation, we know that a bee’s life is made up of multiple bee-days. To ask about bee-lives with similar patterns and changes of activity as a bee ages, we filter and transform the data, and perform PCA on the behahvioral metrics of each bee over time.

The tensor *B*_*αtk*_ is used to represent bee-lives, where *α* is for individual bees, *t* is an index over the days in the bee’s life, which goes from 0 to *l*_*α*_, where *l*_*α*_ is the total number of days the bee lived, and *k =* 1..13 is an index over the component values of **M**. To analyze how different one bee’s life is from another’s, we must consider that all bees did not live for the same number of days. Because of this, we use a parameter *A*_*max*_ *=* 25 for the maximum age used in the bee-life analysis. Because some bees did not live a total of *A*_*max*_ days, and even if a bee was alive there could be some days where it was not detected by the tracking system, we only include bees for the lifetime analysis that had *D*_*min*_ *=* 10 days or more in the bee-day data matrix. With these criteria, and also only keeping bees from cohort G onward, i.e. bees with birthdates within the observation period, we include 2042 bees in the bee-life analysis. We note that *A*_*max*_ and *D*_*min*_ are analysis parameters and quantitatively affect results, although we found that different values of these parameters lead to qualitatively similar interpretations in the differences among bee-lives. We used averages to fill in values of the bee-life matrix for the purposes of PCA and clustering. Let *h*_*tk*_ *=* ⟨ *B*_*αtk*_ ⟩_*α*_, where the notation ⟨ · ⟩ _*α*_ represents an average over the index *α*, denote the average behavioral metrics for each day of the lives of bees that were observed. For a bee that was dead or not observed on day *t* of its life, we fill these values by setting B_*αtk*_|_bee *α* dead or notobserved on day *t*_*= h*_*tk*_. We use *h*_*tk*_ instead of zeros to fill values for the bee-life distance metric, because although the column average of **M** is zero, the average conditional on the age of the bee is nonzero, and therefore filling with zeros would bias the results. After this filtering and processing, we use the bee-life matrix *B*_*αtk*_ as input to PCA and clustering, to obtain the results shown in Figure 6.

We obtain that PCA 1 explains 14.9% of the total variance, PCA 2 explains 7.5% of the total variance, further components explain a smaller fraction of the total variance (Figure S7). Note that because the input is high-dimensional, with *A*_*max*_ * 13 = 325 columns, the fraction of the variance explained by any single mode is relatively small, with an average at 0.31%, and therefore the first two modes represent strong patterns in the data because they are very high above this average variance fraction.

We use *B*_*αtk*_ as input to Ward hierarchical clustering obtain the clusters shown in Figure 6, where the results for a 5 cluster division are highlighted. More clusters can be used to used to describe trends in more detail. Using an 8-cluster division further divides the groupings; however, the differences are still mainly along life PCA components 1 and 2 (Figures S9, S10). From the 5 cluster division to the 8 cluster division, we can describe the further divisions as follows: Cluster 1 (high dispersion - positive PCA 1 embedding values) → clusters 2 & 3, which both have high and positive, although slightly different median projection values, onto PCA component 1, and but differ in the projection on PCA component 2, reflecting differences in nest use with age between clusters (Figures S9). Cluster 4 (early to forage - positive PCA 2 embedding values) → clusters 5 & 6, which further differ along PCA component 1. Cluster 5 (average values) → clusters 7 & 8, which differ predominantly along PCA 1. With this we see that although with 8 clusters there are more groupings, the differences among clusters with the 8-cluster division still reflect the first two dominant axes of variation (Figures S9, S10).

