## Supplemental Figures for "The dominant axes of lifetime behavioral variation in honey bees"

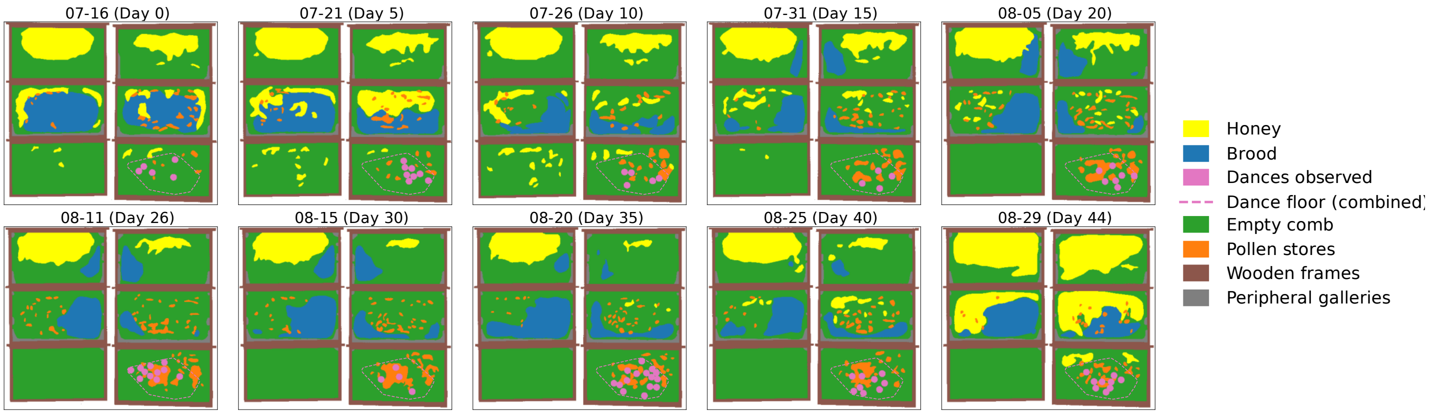


**Figure S1. Comb contents over the observation period.** Figure shows the nest contents and tracings. The pink dashed line in each shows the "combined dance floor", which is defined as a convex hull that contains the locations of all observed dances. The relative locations of the nest contents remained stable throughout the experiment, with honey stored at the top of the nest, brood reared in the center, and a dance floor at the bottom of the nest, near the nest entrance.

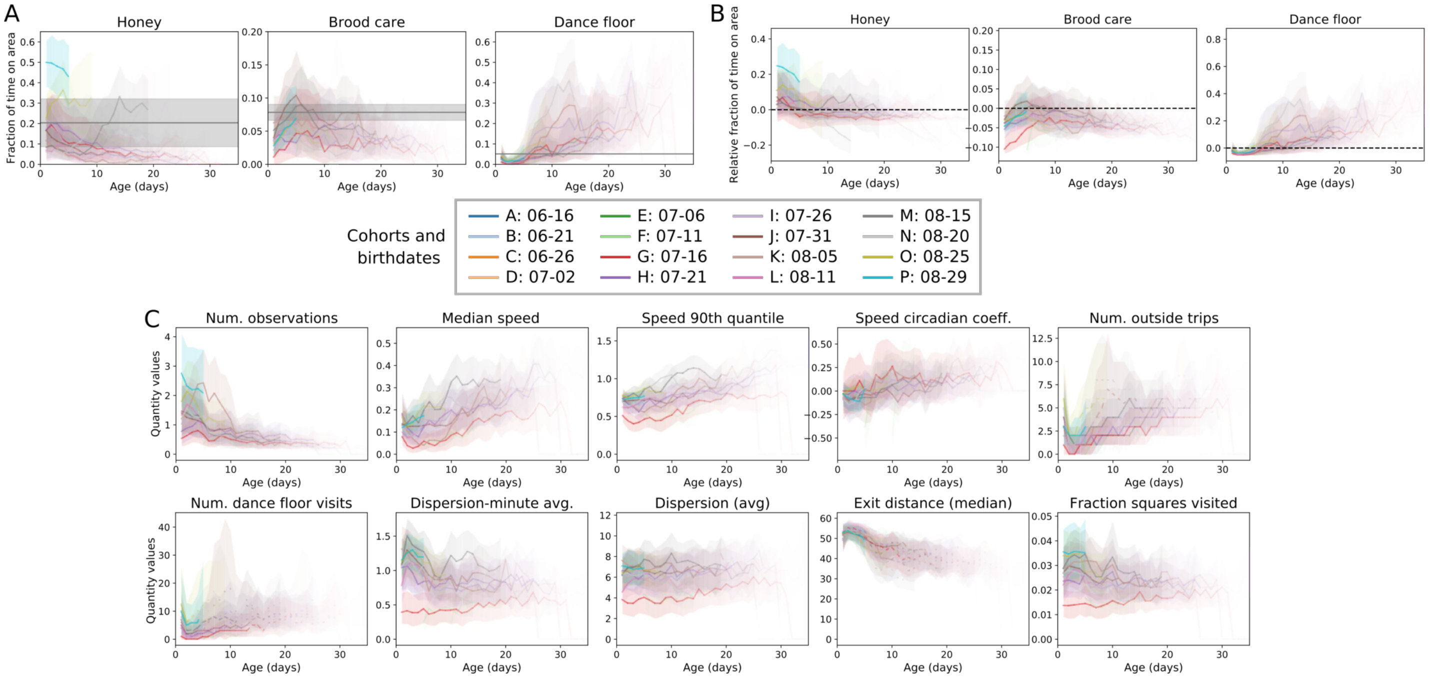


**Figure S2. Substrate and other quantities with age.** See also Figure 2. Cohorts are indicated by the different colors for each plot. (A) Fraction of time spent on honey and brood. The gray line and shaded area shows the mean and standard deviation of the amount of honey or brood in the nest over time. (B) Fraction of time on honey and brood relative to nest contents, determined by subtracting the average contents of the nest for a given day. (C) Other behavioral metrics with age.

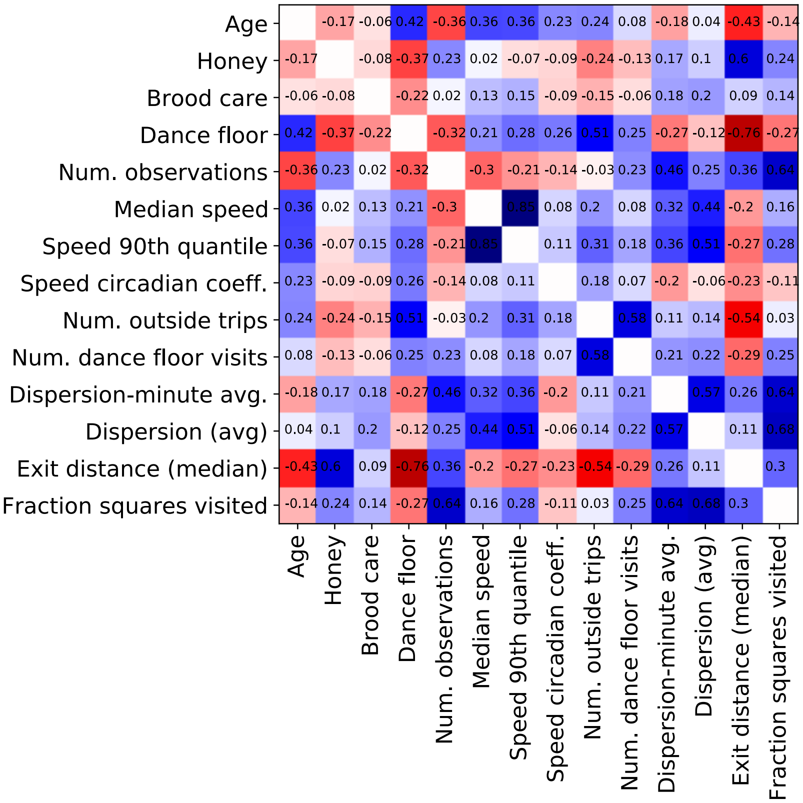


**Figure S3. Correlation between age, substrate use, and other behavioral metrics.** Pearson (ranked) correlation coefficient shown, for all behavioral metrics as well as age. Blue indicates a positive correlation, and red indicates a negative correlation; the values of the correlation coefficient are shown for each pair of quantities.

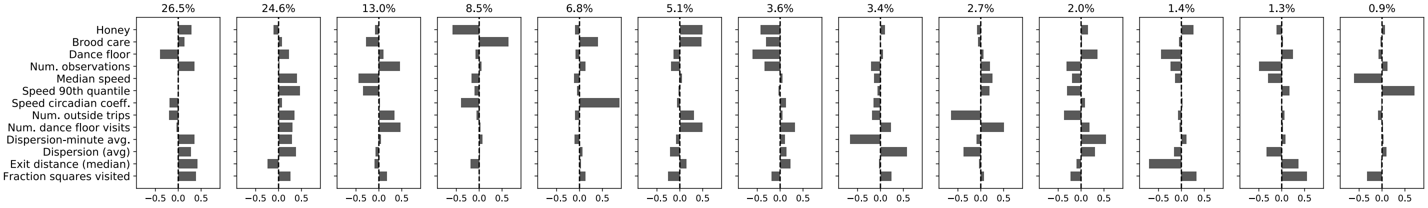


**Figure S4. All bee-day PCA components.** See also Figure 3. The percent variation explained by each component is shown at the top of each.

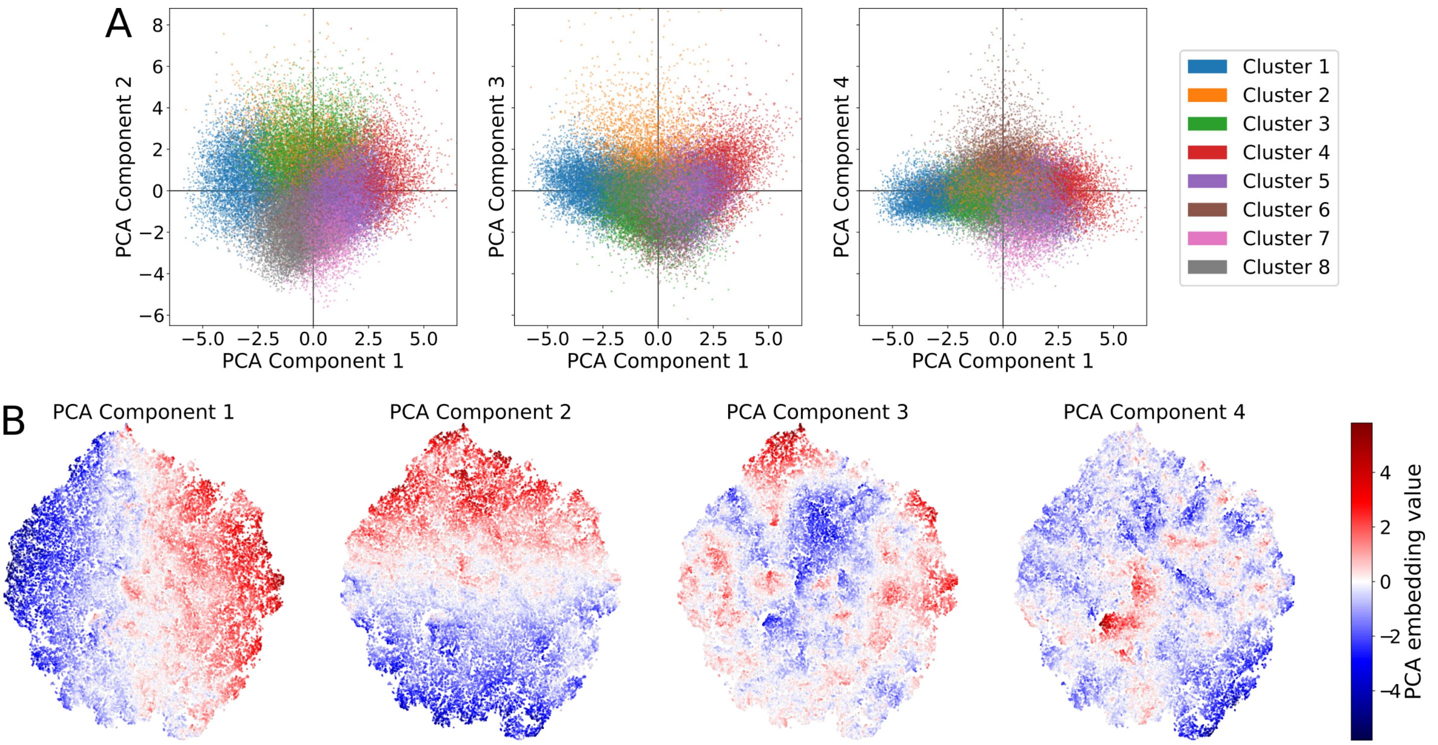


**Figure S5. PCA embeddings of bee-day data.** See also Figures 3, 4, S4. (A) The PCA component projections of the bee-day data, plotting projection values along the first PCA component versus the 2nd, 3rd, and 4th components. See Figure S4 for the PCA component vectors. Colors correspond to the different clusters highlighted in Figure 4. (B) t-SNE embedding of the bee-day data, colored by the projection values along each PCA component dimension. The t-SNE is initialized with the first two PCA component projections, and therefore the global structure of the t-SNE embeddings aligns with these projections.

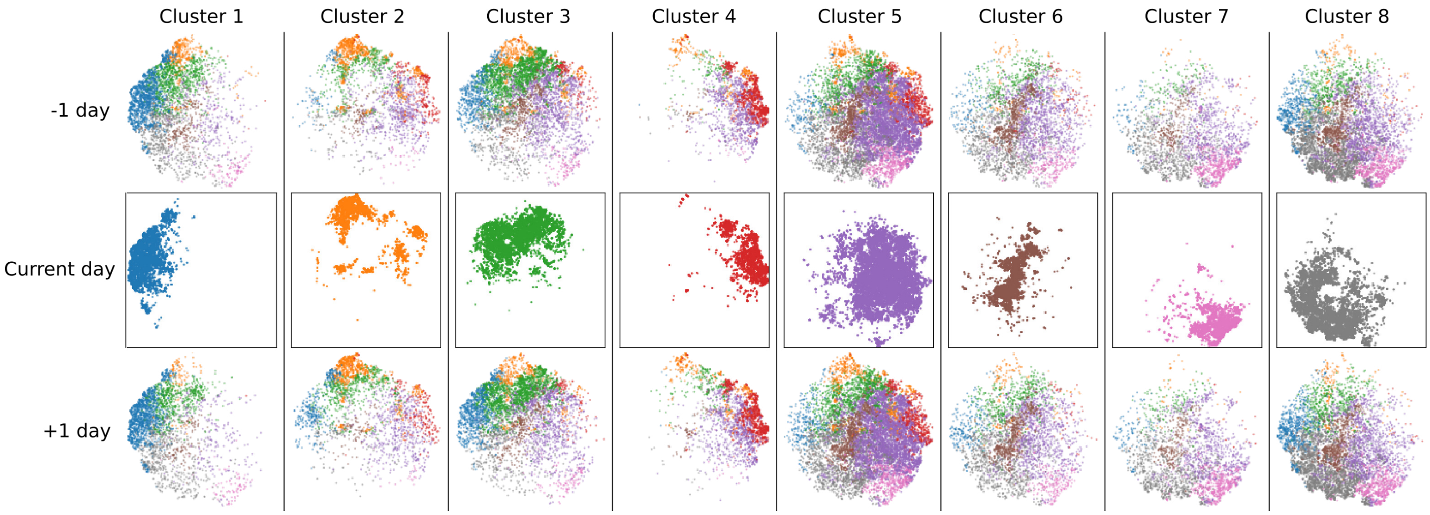


**Figure S6. Embeddings on subsequent days, conditional on behavioral cluster**. The t-SNE embedding of bee-days colored by cluster, a day before or after a given bee had activity within one of the 8 clusters shown in Figure 4. Cluster definitions and colors are the same as Figure 4.

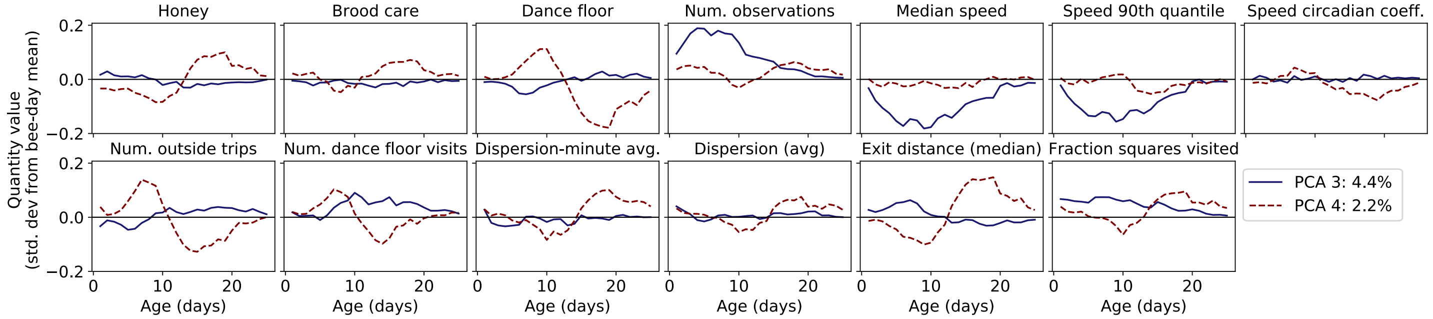


**Figure S7. Further PCA modes from the bee-life decomposition.** See also Figure 6. Life-PCA 3 accounts for 4.4% of the total variance, and PCA 4 for 2.2%. (From Figure 6, the variance fractions for PCA 1 and 2 are 14.9% and 7.5%, respectively). Note that these fractions of the variance are generally small due to the high dimensionality of the input (total number of columns for PCA is $A_{max} * 13 = 325$).

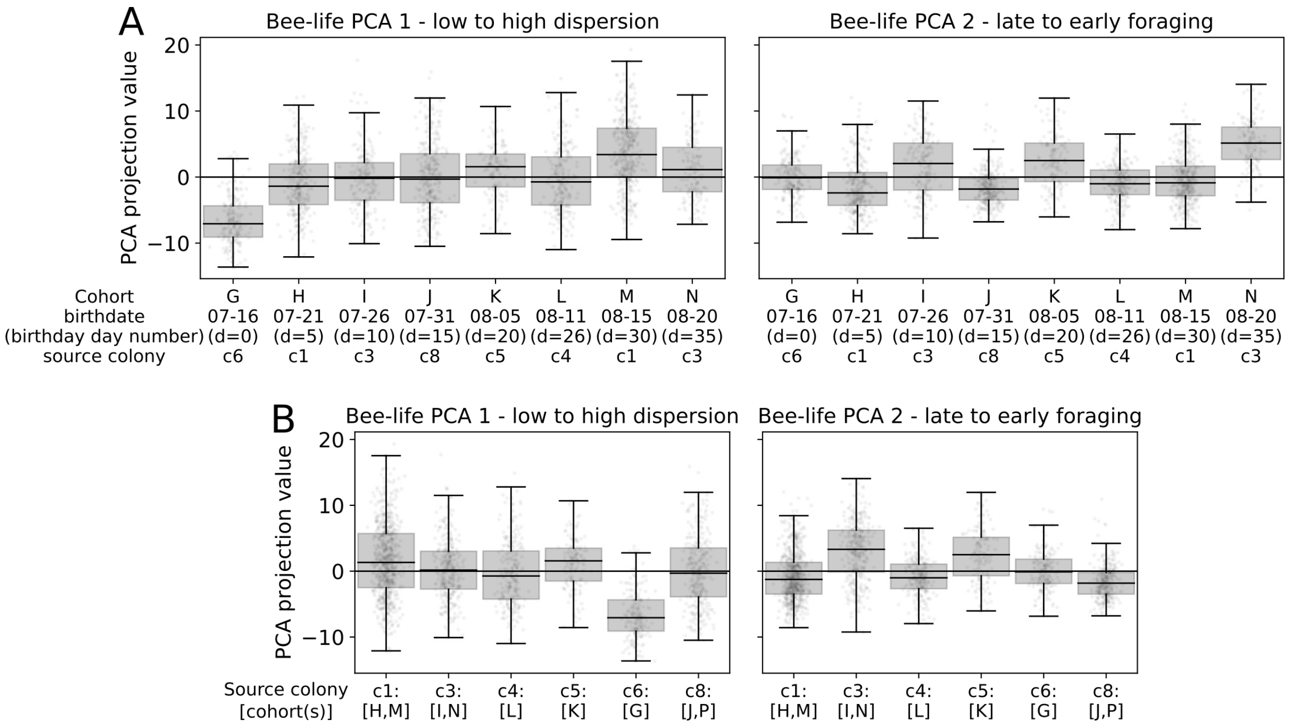


**Figure S8. Cohort and source colony distributions of lifetime behavior.** The distribution of life-PCA embeddings for bees that were included in the lifetime analysis. The bee-life PCA modes 1 and 2 are shown in Figure 6. (A) Per-cohort distributions of lifetime PCA 1 and 2 projections. Note that only cohorts from G onward are included in the lifetime analysis, because these bees have birthdates within the observation period. Cohorts are sorted chronologically, with birthdate and associated source colony shown in the label. (B) Per-source colony distributions of lifetime PCA 1 and 2. Associated cohorts are listed in the label.

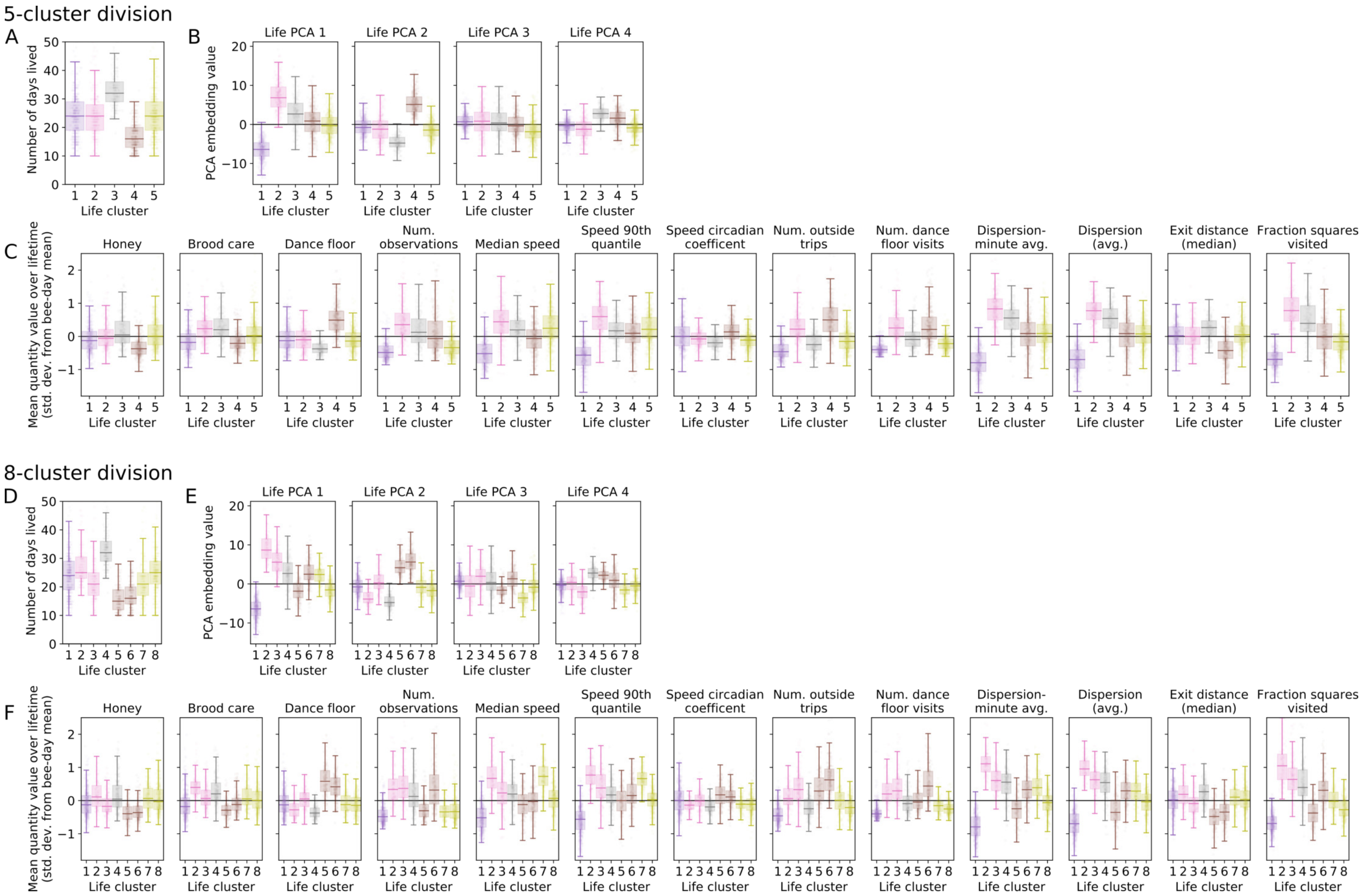


**Figure S9. Life PCA projections and average behavioral metrics, grouped by bee-life cluster.** (A,D) Distributions of the number of days lived for bees in each life cluster. (A) is repeated here from Figure 6E for ease of comparison. (B,E) Bee-life PCA embedding values, obtained by projecting lifetime behavioral metrics onto the lifetime PCA decomposition shown in Figures 6A and S7. (C,F) The distributrion of the lifetime average of each behavioral metric, grouped by bee-life cluster. Results are shown for a division into (A-C) 5 clusters, as highlighted in the main text in Figure 6), and (D-F) 8 clusters. The colors from the 8 cluster result reflect the branching divisions from the 5 cluster grouping: Cluster 2 from the 5 cluster grouping divides to become clusters 2&3 in the 8-cluster division, etc.

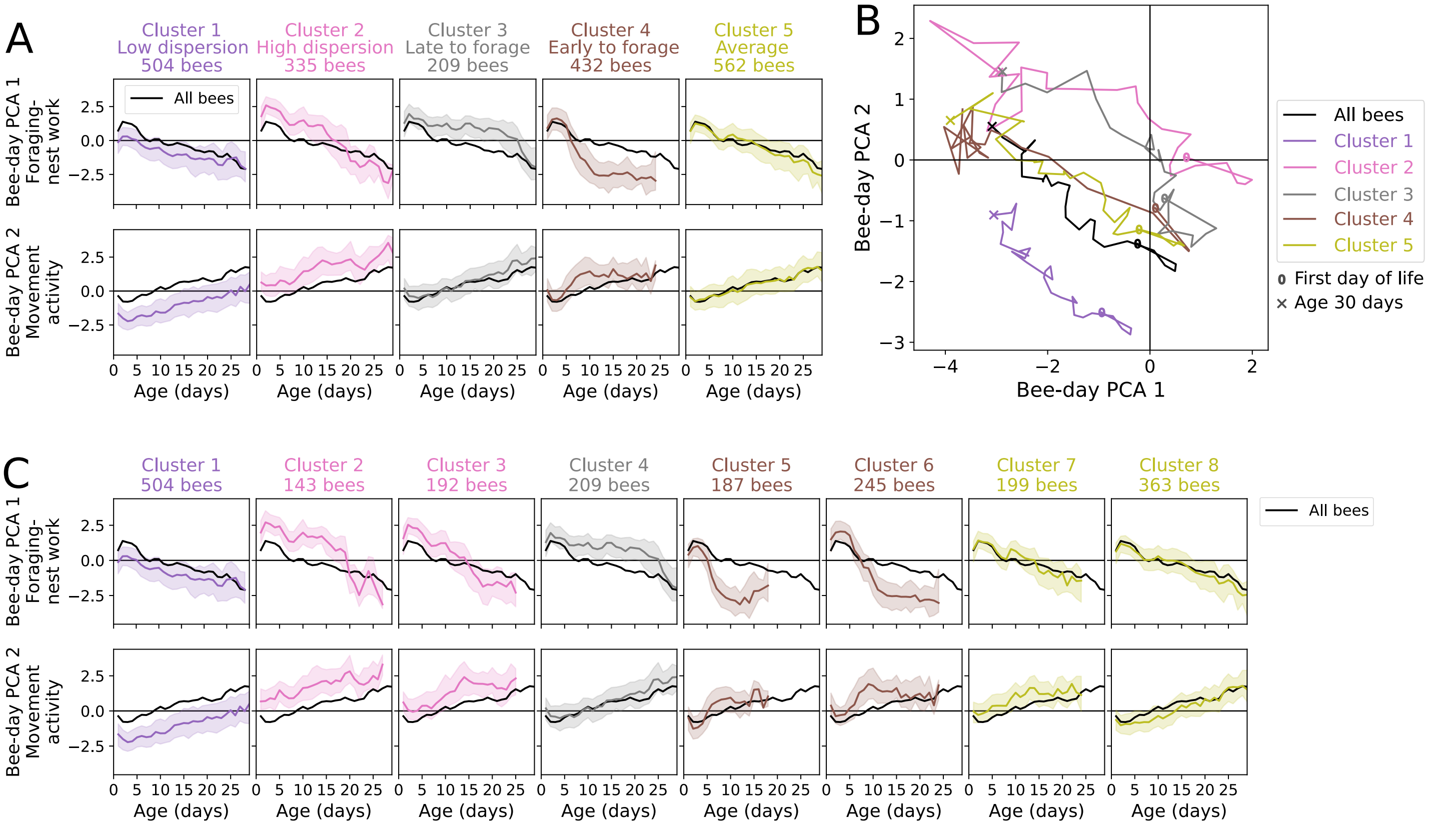


**Figure S10. Distributions of bee-day PCA component values with age, grouped by bee-life cluster.** (A,C) The median values of each bee-day PCA component projection (solid colored line) along with the inter-quartile range (shaded area), for bees of a certain age in each lifetime behavioral cluster, compared to the median value of that PCA component projection for all bees as a function of age (black line). Results are shown for a division into (A) 5 clusters, as highlighted in the main text in Figure 6, and (C) 8 clusters. As in Figure S9, the colors from the 8 cluster result reflect the branching divisions from the 5 cluster grouping: Cluster 2 from the 5 cluster grouping divides to become clusters 2&3 in the 8-cluster division, etc. (B) The median values of the dominant two bee-day PCA component projections, grouped by life cluster, and plotted with age. The '0' denotes the median values at the start of the bees life, and the 'X' denotes values at age 30.

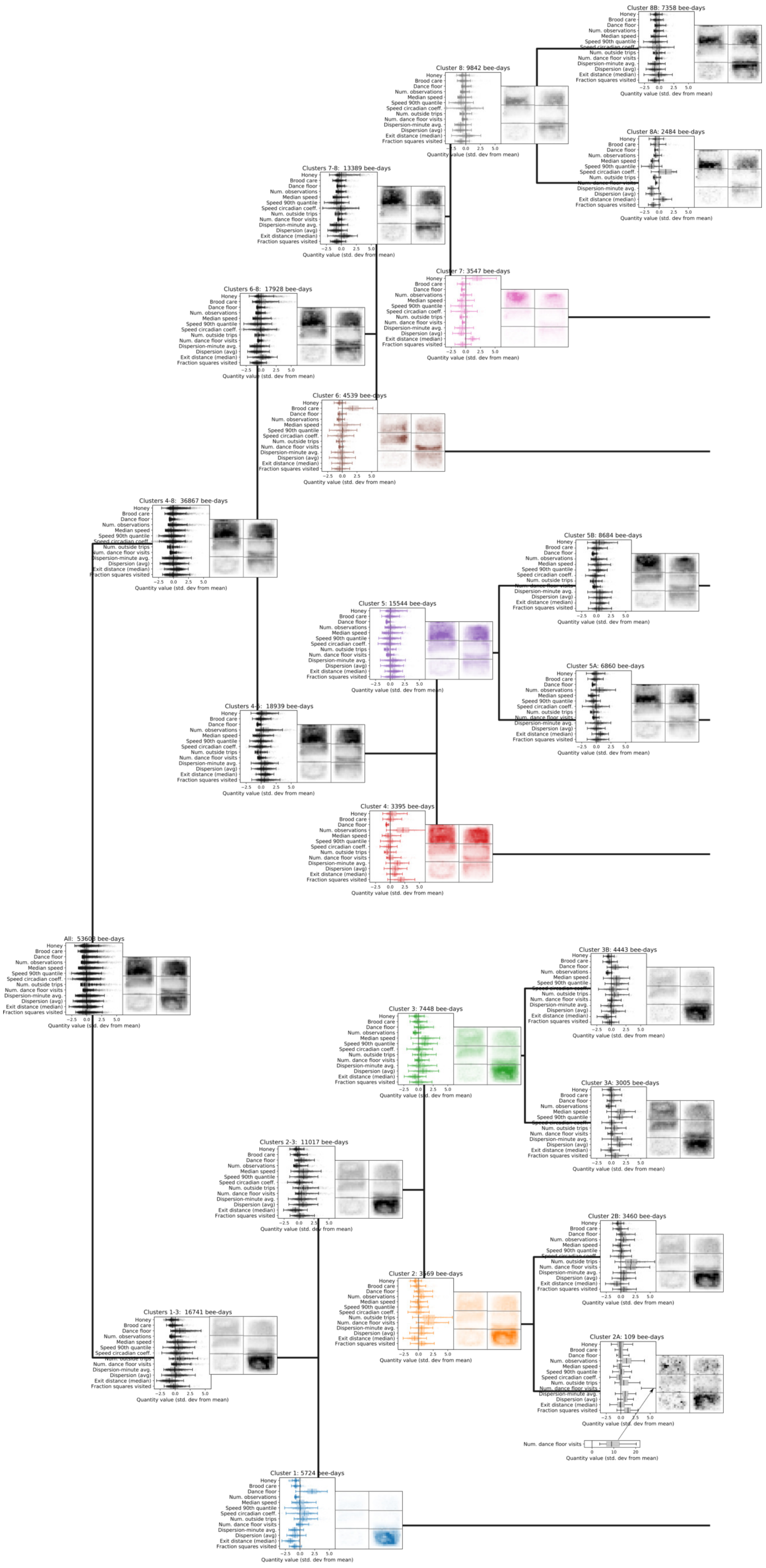


**Figure S11. Dendogram and feature diagram**. See also Figure 4. Quantity distributions are shown along with averaged histograms for bee-days in each cluster.
